# Isolation and Purification of Lipoarabinomannan from Urine of Adults with Active Tuberculosis

**DOI:** 10.1101/2021.03.24.436904

**Authors:** Jason L. Cantera, Andrew A. Rashid, Lorraine L. Lillis, Roger B. Peck, Paul K. Drain, Abraham Pinter, Masanori Kawasaki, Emmanuel Moreau, David S. Boyle

## Abstract

Lipoarabinomannan (LAM) is a cell wall component of *Mycobacterium tuberculosis* that is excreted in the urine of persons with active tuberculosis (TB). Limited diagnostic sensitivity of LAM immunoassays has been due to selecting antibodies against LAM derived from *in vitro* cultured *M. tuberculosis*, rather than LAM purified from *in vivo* clinical urine specimens. Urinary LAM (uLAM) is critical to enable the development of and/or screening of novel uLAM-specific antibodies but is typically dilute and in heterogeneous mixtures with other urine components. We used physical, enzymatic, and chemical processes for the scaled isolation and purification of uLAM. The purified material may then be used to develop more sensitive uLAM diagnostic tests for active TB disease.

## Introduction

Tuberculosis (TB) is a treatable infectious respiratory disease that causes 10 million new active cases each year, and over 1.4 million deaths in 2019 (36). TB is caused by a bacterium, *Mycobacterium tuberculosis*, which typically infects the lungs, but can also cause extra-pulmonary infections. All mycobacterial species including *M. tuberculosis* produce lipoarabinomannan (LAM), a heterogeneous lipopolysaccharide that is a major component of the mycobacterial cell wall (18). LAM has a glycerophospholipid terminus, which non-covalently anchors the molecule within the inner cell membrane, and attaches to a common mannan domain (6). The number of and presence of sugars that cap the terminal residue of the arabinan side chains discriminate different species of mycobacteria (34). The arabinan side chains of the fast-growing species (e.g., *M. smegmatis*) are uncapped or have inositol phosphate caps. Conversely, slower growing species (e.g., *M. tuberculosis, M. leprae*) have caps of one to three units of α (1→2)-linked mannopyranose (Man*p*) (5). A further *M. tuberculosis-specific* modification is the capping of the terminal Man*p* residue with 5-deoxy-5-methylthio-xylofuranose (33). LAM’s average molecular weight is 17.4 kilodaltons although multiple species are formed depending on the variety of modifications including the size, branching, acylation and phosphorylation of the mannan and arabinan components (6,18).

Soluble LAM is actively secreted from bacteria and infected macrophages and acts as a virulence factor (2,8,23). Soluble LAM is relatively abundant, produced in all types of TB disease (pulmonary and extrapulmonary) and ultimately is filtered by the glomeruli and excreted in urine, and therefore making it a compelling biomarker for use in non-invasive rapid diagnostic tests for TB (7,11,22,27). Several rapid diagnostic tests based on urinary antigen immunoassays targeting urinary LAM (uLAM) have been described, but none are sensitive enough to reliably detect uLAM for active TB (3,20). Efforts are ongoing to improve the sensitivity of rapid uLAM tests via additive methods such as a larger sample volume (3), sampling first pass morning collection versus later urine sampling (13), enrichment of LAM prior to testing (26) or with pretreatment prior to testing (12,17,25).

Most of the anti-LAM monoclonal antibodies described in the literature use immunogens purified from *in vitro* cultured *M. tuberculosis* cells and/or using culture-derived LAM for screening the candidate clones derived (9, 14–16,19,28). While this is convenient in terms of producing sufficient quantity and high-quality material for these purposes, *in vivo* cultured *M. tuberculosis* may produce LAM and/or uLAM with significant structural differences (10). Therefore, clinical performance may be compromised in part by key epitopes in these derivatives of LAM not being recognized (10,21,29). In this study, we describe a series of chemical, enzymatic, and physical steps to enrich and purify soluble LAM from the urine of patients confirmed with TB to demonstrate that it is possible to do this at scale and create sufficient amounts of purified LAM which may further support early development and performance verification of uLAM immunoassays in place of LAM from *in vitro* cultured *M. tuberculosis*.

## Materials and Methods

### Clinical specimens

Urine specimens from 40 TB positive patients were obtained from the FIND and WHO Tuberculosis Specimen Bank (Geneva, Switzerland) and the uLAM concentration for each confirmed via a reference immunoassay for TB LAM and uLAM (4,31). A 500 mL volume of pooled urine was prepared with 1 mL aliquots from TB positive specimens (40 mL total) and added to 460 mL of pooled urine collected from healthy individuals (BioIVT, Westbury, NY, USA). The contrived urine was stored overnight at 4 °C prior to processing.

### Immunoassay measurement of LAM concentration

Purified LAM derived from *in vitro* cultured *M. tuberculosis* strain Aoyama-B was procured from Nacalai USA, Inc. (San Diego, CA, USA). Monoclonal anti-LAM antibodies, including A194-01 (Rutgers University, New Brunswick, NJ, USA), FIND 28 (FIND, Switzerland), and S4-20 (Otsuka Pharmaceutical, Japan), were used to prepare sandwich immunoassays to detect LAM in a biplexed format as previously described (31). The immunoassays were performed using U-PLEX 96-well plates and analyzed using a proprietary plate reader (MESO QuickPlex SQ 120, Meso Scale Diagnostics, LLC, Rockville, MA, USA), and results analyzed using Discovery Workbench 4.0 (Meso Scale Diagnostics).

### Purification of urinary LAM

The purification scheme used in this work is outlined in Figure 1. Where feasible, the collected and discarded material from each processing step was assessed for uLAM concentration via the MSD LAM immunoassays. A precursory step (Step 1) was employed to remove gross particulates, such as bacterial cells (including *M. tuberculosis*) and cellular debris from the 500 mL of urine by centrifugation at 1,200 rpm for 15 min. The resulting supernatant was further clarified by micro-filtration using a 0.2 μm filter (Sarstedt Inc, Newton, NC, USA) to remove microparticulates. This filtered volume was considered the starting material for the enrichment and purification steps. The filtered urine was initially concentrated into approximately 50 mL using a Vivaflow 200 diafiltration instrument (Sartorius, Göttingen, Germany) using a flow rate 80 of mL/min, and the concentrate then washed *in situ* with ice-cold phosphate buffer saline (PBS) pH 7.4 at 5 times the concentrate volume (Step 2). The concentrate was then treated with proteinase K (Fisher BioReagents, Waltham, MA, USA) to 200 μg/mL final concentration for 1 hour at 55 °C (Step 3). Protease activity was inactivated by heating the reaction to 100 °C for 10 min.

**Figure 1.**
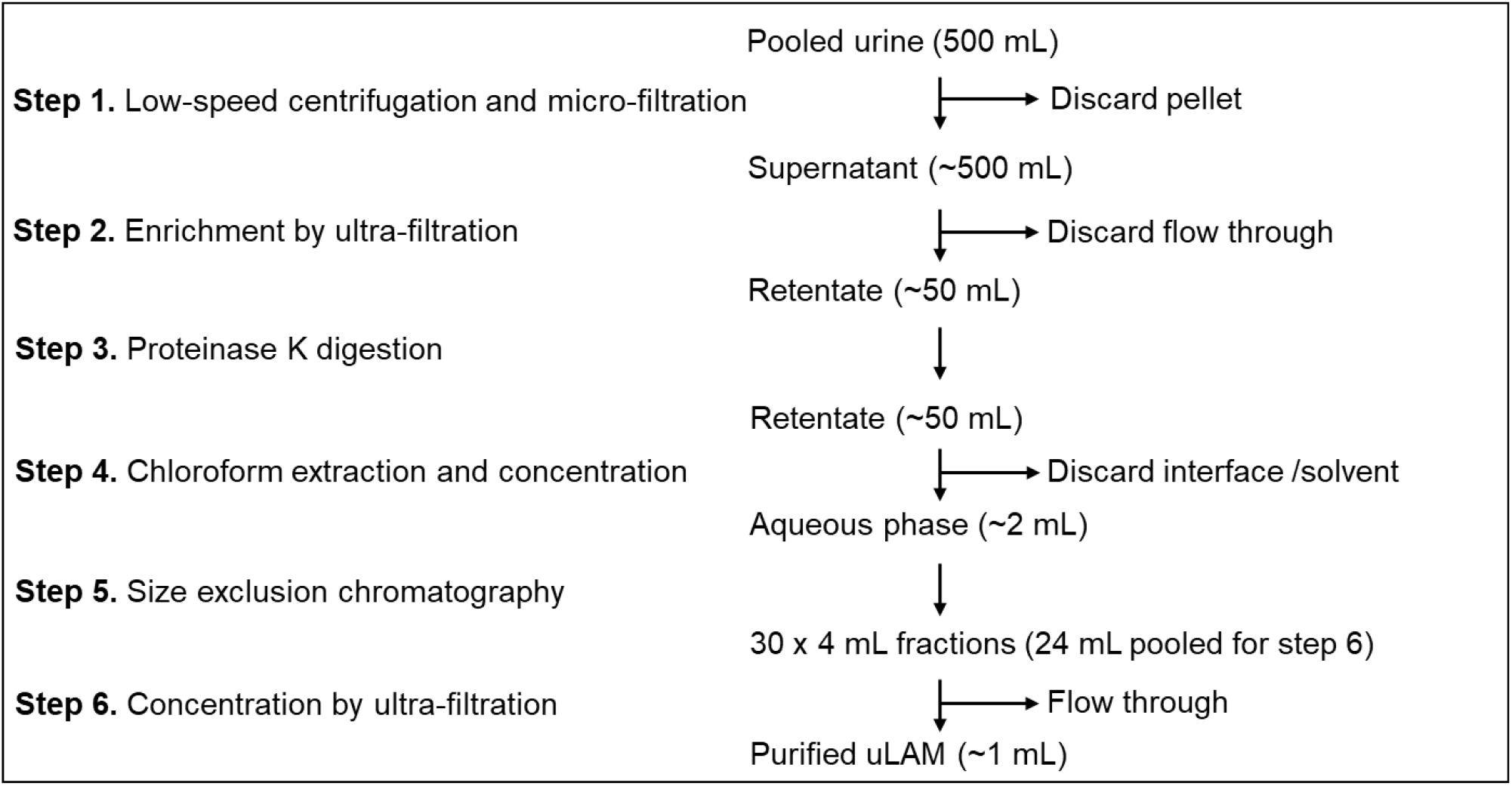
A flow diagram depicting the sequential processing methods (Steps) used to purify and concentrate uLAM from 500 mL of pooled urine.

After proteinase K digestion, the digest was solvent-extracted in an equal volume of chloroform (Millipore-Sigma, St. Louis, MO, USA). The sample was mixed by vortexing for 30 seconds and then centrifuged at 4,000 rpm for 10 minutes. The aqueous phase was collected, and solvent phase was re-extracted using PBS (v/v 0.5/1.0). The two aqueous phases were combined and then concentrated to a final volume of 2 mL by ultra-filtration using a 10 kDa MWCO Ultra-15 centrifugal filter (Millipore-Sigma).

The concentrated material was then subjected to size exclusion chromatography (Step 5) using PBS-equilibrated HiPrep^™^ Sephacryl S-100 HR column (100 x 2.5 cm, Millipore-Sigma) in an AKTA start (Cytiva, Marlborough, MA, USA) with a flow rate of 0.5 mL/min. Thirty 4-mL fractions were collected and assayed for the presence of uLAM using the reference LAM immunoassay. Fractions that strongly reacted with both antibody pairs were pooled and concentrated using a 10 kDa Ultra-15 centrifugal filter (Millipore-Sigma).

The total LAM concentration from each step was measured by the two optimal immunoassays described by Sigal *et al*. (31). The two assays were combined into a biplexed assay employing two antibody pairs as capture and detector: FIND28/ A194-01 (FA) and S4-20/ A194-01 (OA), which recognize different epitopes in the LAM structure and therefore provide different estimates of LAM concentration. The depletion of both proteins and carbohydrate levels in the urine was also measured after each processing step (Figure 1). The total protein concentration was measured using the BCA method (ThermoFisher Scientific, Waltham, MA. USA). Total carbohydrate measurement used a phenol-sulfuric acid assay (24). Two different standards were used to measure the total carbohydrate concentrations, one using glucose (Sigma-Aldrich) and the other purified Aoyama-B LAM from *in vitro* culture.

## Results and Discussion

The efficiency of purification of LAM from urine was directly assessed for LAM and for the depletion of both total protein and carbohydrates from the samples (Table 1). No detectable LAM was measured in the pellet produced from an initial centrifugation step. The stock urine volume was also passed through a 0.2 μm filter to remove small particulates from the sample (Step 1). As noted earlier, the total uLAM concentration measured in the samples differs depending on which antibody pair is used (31). The total amount of LAM in the 500 mL starting material from Step 1 was estimated to be 486.9 ng or 102.8 ng using FA and OA assays, respectively. The total protein was 14.35 mg and for carbohydrates, the absolute concentrations were also different at 484.3 μg and 258.7 μg when using LAM or glucose, respectively, for the standard.

**Table 1.** The sequential processing steps used to enrich uLAM, with measurement of uLAM quantities via two immunoassays. The total proteins and carbohydrates in each derivative were also measured, the latter using one standard of glucose and another using LAM purified from TB culture. The amount and relative percentile to the input amounts in step 1 are shown.

| Step | Technique | uLAM |  |  |  | Protein |  | Carbohydrate |  |  |  |
| --- | --- | --- | --- | --- | --- | --- | --- | --- | --- | --- | --- |
|  |  | FA |  | OA |  |  |  | LAM std |  | Glucose std |  |
| | | ng* | % | ng* | % | $\mu\text{g}^*$ | % | $\mu\text{g}^*$ | % | $\mu\text{g}^*$ | % |
| 1 | Microfiltration | 486.9 | - | 102.8 | - | 14349.6 | - | 484.3 | - | 258.7 | - |
| 2 | Ultrafiltration | 403.9 | 83.0 | 73.1 | 71.0 | 69.9 | 0.5 | 14.0 | 2.9 | 8.0 | 3.1 |
|  |  | (26.6) | 5.5 | (24.6) | 23.9 | (13409) | 93.4 | (454.3) | 93.8 | (249.4) | 96.4 |
| 3 | Proteinase K | 463.6 | 95.2 | 74.1 | 72.1 | 76.9 | 0.5 | 15.6 | 3.2 | 8.9 | 3.5 |
| 4 | Chloroform extraction | 309.0 | 63.5 | 47.1 | 45.8 | 2.8 | 0.02 | 8.7 | 1.8 | 5.3 | 2.0 |
| 5, 6 | Size exclusion chromatography followed by ultrafiltration | 250.4 | 51.4 | 32.1 | 31.2 | 2.3 | 0.02 | 4.1 | 0.9 | 1.3 | 0.5 |

\*Quantities are normalized per mL of material/ resulting fractions in each step; in parenthesis are quantities in the filtrate after ultrafiltration.

In Step 2 the material was passed through a 10 kDa MWCO ultra-filtration system to reduce the overall sample volume, enrich soluble uLAM in the retentate, remove some of the smaller soluble molecules from the sample, and allow buffer exchange of solutes. This treatment resulted in 83.0% and 71.0% of uLAM being retained while filtration removed 93.4% of the total proteins and likely all the carbohydrates with either standard indicating 93.8 to 96.4% of the carbohydrates were removed. About 5.5% and 23.9% uLAM were in the filtrate. It is possible that LAM, with an average MW of 17 kDa, was partially fragmented in urine in sizes small enough to pass through the 10 kDa filter. The slight loss of uLAM was also likely due to non-specific binding to the filter or the uLAM being complexed with other larger molecules that tightly bound to the filter membrane. The enrichment of uLAM via ultrafiltration reduced the initial volume of ~500 mL urine to ~50 mL which enabled proteinase K digestion of the remaining soluble proteins co-enriched after the concentration (1). This treatment slightly altered the LAM measurement with FA recording 60 ng more than in the previous step (2). This is possibly the result of more epitope being revealed after proteolysis, a feature previously noted with proteinase K treatment of urinary LAM (1). Both LAM measurements were equivalent to the pretreatment values and this was to be expected since only proteins were affected by this step. Similarly, the protease K treatment did not alter the total protein concentration as the proteinaceous material remained, albeit most of it as digested peptides.

The removal of these peptides by solvent extraction in Step 4 resulted in the greatest single loss of LAM in the entire process despite backwashing the solvent phase. The losses of LAM were 25% irrespective of the LAM assay used. Some LAM may have been irrecoverable as trapped in the matrix of insoluble denatured peptides that formed when the sample was mixed with chloroform. The effect of solvent extraction on the total protein concentration was very significant in that with the removal of a further 74.1 mg/mL, only 0.02% of the original amount of protein remained. The use of size exclusion chromatography was considered as a polishing step by which more soluble material could be removed from the sample with residual larger molecules washed out. The distribution of uLAM in the fractions collected can be seen in figure 2. Fractions 10 through 17 were observed to contain uLAM and the peak using OA was much less pronounced than that for FA. The LAM containing fractions were sequentially concentrated to create a final pooled fraction of 1 mL (Figure 2). The carbohydrate levels in this sample dropped to 0.5% of the initial amount although the significance of this to the previous measurement is dubious as this is likely within range of intra-assay variation, especially with samples that are towards the assay’s lower limit of detection. Step 5 did have a negative effect on the total uLAM concentration with similar percentile loss of uLAM at 12.1% and 14.6% when measured by FA and OA, respectively.

**Figure 2.**
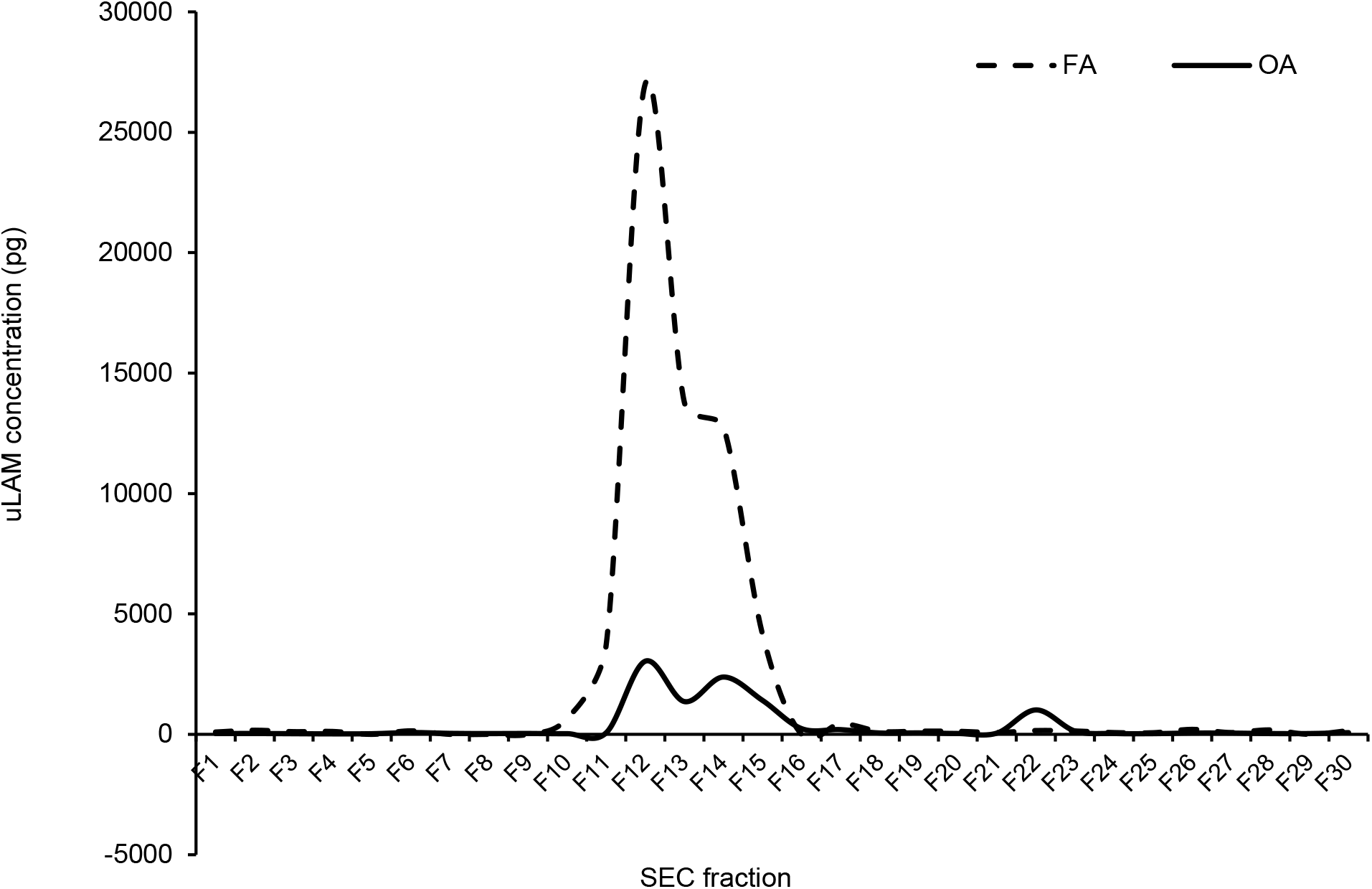
uLAM immunoassay analysis of the fractions collected by SEC using FA (dashed line) and OA (solid line) assays.

In summary, we first assessed the performance of a variety of individual methods that when combined can effectively enrich and purify uLAM from the urine of infected TB patients with a reasonable yield of product. The methods and equipment necessary are inexpensive and familiar to the typical modern molecular laboratory and compromise a mixture of basic physical processes such a centrifugation and filtration combined with enzymatic and chemical treatments. The use of LAM-specific affinity immuno-chromatographic methods was deliberately omitted to reduce the bias with which specific LAM moieties or derivatives were captured. An alternative non immuno-affinity method could be to use recombinant human mannose-binding lectin which has been shown to interact with mannan-capped LAM (32). However, this is expensive if used at scale and in addition, with the heterogeneous nature of uLAM it is possible that a subpopulation of LAM may no longer be MAN-capped. Of all the steps, solvent extraction was the least efficient in terms of retaining uLAM, likely due to loss of LAM in the insoluble peptide aggregates. This step did however achieve the goal of removing most of the residual proteinaceous material after ultrafiltration and proteinase K digestion.

We demonstrated a purification efficiency of up to 51.4% from a deliberately contrived urine sample at medium scale volume (500 mL) to significantly increase the amounts of uLAM that can be potentially purified. In our laboratory we have observed patient samples with uLAM concentrations as high as 82.5 and 23.4 ng/mL using FA and OA assays, respectively. We envisage that urine samples with grade 4 scores from the Alere Determine TB LAM assay could be collected in large volumes and pooled and processed using the method described to yield much larger amounts of uLAM. This would further qualify the development and verification of new or existing TB LAM diagnostic assays to better meet the clinical performance ranges required to meet the specifications of current target product profile for either triage tools or rapid diagnostic tests to inform on TB infection (35).

Our study has several limitations. Firstly, the amount of purified uLAM was very low and insufficient for typical molecular analysis of LAM via SDS-PAGE and western blots that would support our claims of increased purity and yield and allow direct comparison to LAM derived from cultured *M. tuberculosis* (16,30). We did not measure the salt concentration of the samples and assume that the dilutions with PBS in Steps 1 and 5 removed many inorganic solutes without providing direct evidence of this. As seen with FA and OA assays, each gives a different estimate of concentration as reflected partly by the specificity of the antibodies used but also in the prevalence of specific epitopes found in uLAM, a likely very heterogeneous mixture of uLAM structures (9). The accuracy of some measurements varies with the purity of the sample increasing and providing less matrix effects, the proteinase K treatment significantly improved the recovered amount via FA, but not for OA and we hypothesize more epitopes were exposed after this treatment, something previously observed with an immunoassay after using proteinase K treatment, albeit with a different antibody pair (1). The uLAM produced by *M. tuberculosis* infection in different hosts and by different TB lineages is yet unknown and so uLAM is very likely to be highly heterogenous when these factors are considered and is in need of further research to understand the effects of these on the performance of rapid diagnostic tests for the presence of LAM in urine.

By offering a basic methodology that can effectively enrich uLAM from large volumes of urine, we are aiming to process much larger volumes of urine with higher uLAM concentrations in order to create stocks that can be used to more accurately inform on LAM antibody development across the development spectrum from the selection of B cells or their derivatives producing uLAM-specific antibodies, to screening new antibodies in order to investigate the binding kinetics and performance at an early stage of development using the direct target versus the cultured surrogate. With better materials to optimize the performance of urinary LAM immunoassays, the target of a rapid triage test to diagnose TB directly from urine is more likely to succeed.

## Acknowledgements

This work was funded via a grant (INV-008079) awarded to DSB by the Bill and Melinda Gates Foundation (https://www.gatesfoundation.org/). The funder did not have any additional role in the study design, data collection and analysis, decision to publish, or preparation of the manuscript. We would like to thank Drs. Delphi Chatterjee and Prithwiraj De and Ms. Anita Amin from Colorado State University for their support in understanding the variance and structural complexity of LAM.

